# Speeded Inference Game: Opening a new chapter in the assessment of error awareness

**DOI:** 10.1101/2022.03.01.481617

**Authors:** Eva Niessen, Jonas Wickert, Martin Schober, Gereon R. Fink, Jutta Stahl, Peter H. Weiss

## Abstract

Influential theories on error processing assume that when we conduct errors adaptive processes are triggered to improve our behaviour and prevent errors in the future. These processes appear to be more effective after participants have detected an error. Therefore, the assessment of error awareness allowing a differential analysis of detected and undetected errors in the context of cognitive control and behavioural adjustments has gained more and more attention in the past decades. A common methodological challenge posed on all studies investigating error detection is that the number of undetected errors is usually relatively low.

Here, we introduce a gamified experimental task that uses an adaptive algorithm to generate a robust and stable amount of errors with a high rate of undetected errors. Further, we were able to identify error types, which interestingly differed in terms of their detection rate. Moreover, the game-like appearance of the novel experimental task led to highly motivated participants. The results of the first study were replicated and extended by a second behavioural study. Notably, in study 2, a change in task design specifically modulated error detection, while these changes did not affect the total error rate.

Potential applications of the open-source code will be discussed. With this newly developed paradigm, we wish to lay the ground for future research to understand better (neural) processes associated with error awareness.

## Introduction

The assessment of error awareness in the context of cognitive control and behavioural adjustments is essential and has gained more and more attention in the past decades. Influential theories on error processing assume that when we conduct errors, adaptive processes are triggered to improve our behaviour and prevent errors in the future (Botvinick et al., 2001; Holroyd & Coles, 2002). These processes occur even when we are not aware of the incorrect action (Hester et al., 2005; Nieuwenhuis et al., 2001), but they appear to be more effective after aware compared to unaware errors (Klein et al., 2007). Commonly used measures of behavioural adaptations after errors are slowing of response times (post-error slowing, PES) or increases in accuracy (post-error increase in accuracy, PIA) compared to correct responses (Danielmeier & Ullsperger, 2011). While PIA represents an immediate performance improvement, the functional significance of PES is still under debate. PES might indicate a meaningful adjustment (e.g., a more cautious, hence slower, response in the trial following an error to increase the chance to stop or change the action successfully; (Endrass et al., 2007; Nieuwenhuis et al., 2001). Alternatively, PES could merely represent a meaningless by-product of the occurred error (e.g., processing the infrequent error results in a delay or disruption of the forthcoming processes resulting in slower responses in the trial following an error; Notebaert et al., 2009; Wessel, 2018).

One methodological challenge posed on all studies investigating error processing is to employ an experimental task that results in many errors in all participants (i.e., a high error rate) without causing a performance breakdown. Too many errors might result in high levels of frustration and participants’ disengagement from the task, or in the disability to detect errors at all. In the past, several methods that increase task difficulty and consequently the error rate have been used (e.g., reduce stimulus contrast to induce perceptual conflict or increase number of distracting stimuli; Schreiber et al., 2012; Somon et al., 2019). To investigate the influence of error awareness on adaptive processes, it is mandatory to distinguish between detected and undetected errors^1^. To this aim, the error rate has to be even larger because detected and undetected errors will be analysed separately. Another problem is that the number of undetected errors is usually very low (Kirschner et al., 2020; Niessen et al., 2017; O’Connell et al., 2007; Rabbitt, 2002), preventing a robust characterisation of the processes leading to undetected errors. Further, a fair and meaningful comparison between processes regarding detected and undetected errors is difficult to achieve if rarely occurring undetected errors are contrasted with more common detected errors. Past attempts to provoke more undetected errors for instance reduced the visibility and processing time of the stimulus (Cohen et al., 2009; Steinhauser & Yeung, 2010), leading to perceptual uncertainty and reduced error detection rates (Endrass et al., 2012). Another approach was to increase task duration, resulting in fatigue and more undetected errors, especially at the end of the experimental session (Harty et al., 2014).

While these procedures indeed resulted in a larger number of undetected errors, it is questionable whether the observed errors could have been adequately detected by the participants in the first place because the experimental manipulations most likely prevented an adequate evidence accumulation (Steinhauser & Yeung, 2012). This would mean that it was generally not possible for participants to become aware of their errors. In addition, the applied methods may have triggered mere guessing of correct responses, at least in some trials. If so, differences between the processing of detected and undetected errors would happen at chance rather than under differential mechanisms under scrutiny.

Usually, electroencephalographic (EEG) analysis require a large number of trials and participants need to be motivated across a longer period to prevent their committed errors (Larson et al., 2014). If errors are not important and significant for them, the motivation to adapt one’s behaviour is usually small (Hajcak et al., 2005). Meaningful errors are coupled with emotional appraisal, and the affective response triggered by errors is assumed to result in more robust adaptive processes (e.g., larger PES; Dignath et al., 2020)). However, undetected errors due to long task durations are likely driven by fatigue and indifference towards errors. Therefore, any difference to errors occurring early in the experiment (which are usually detected), could be confounded by secondary processes (e.g., motivation and alertness/ vigilance) and again would not be based on differences in the cognitive processes *per se*.

### Objectives

Here, we open a new chapter in the assessment of error awareness by presenting a new paradigm that aims to overcome the described methodological limitations. The benefits and characteristics of the paradigm will be described in detail, making it easy for interested readers to use and adjust the open-source code of the paradigm. It consists of a combination of commonly used and novel features (e.g., adaptive algorithm for response deadline, hierarchical rules), and the appearance and choice of stimuli make the paradigm appear game-like. The game-like character should lead to high levels of motivation without compromising the strict control of the experimental task. Furthermore, the ecological validity was expected to be higher compared to classical conflict tasks such as the Go/Nogo or flanker task.

The main goal was to establish the new paradigm and inspect the resulting behavioural pattern, primarily focussing on the amount of detected and undetected errors provoked by the paradigm. In Study 1, we showed that already after a short time, a robust and stable error rate with a high number of undetected errors could be found. The behavioural results of study 1 were replicated and extended by study 2. Further, study 2 exemplifies how easy it can be to manipulate one feature of the task (i.e., rule frequency) to specifically modulate one outcome variable of interest (i.e., error detection rate). The second aim of study 1 was to investigate whether the increased statistical power due to the high rates of detected *and* undetected errors help to inform a recently proposed theory on error processing and behavioural adjustments after errors (Wessel, 2018).

## Method

The paradigm and the experimental parameters of the two studies were identical with one exception (i.e., rule frequency, see below). Thus, the following descriptions apply to both studies.

### Participants

We recruited young, healthy, right-handed (Edinburgh Handedness Inventory, EHI; Oldfield, 1971) participants without any history of neurological or psychiatric diseases. Task instructions and questionnaires were provided in German or English (i.e., participants had to be fluent in German or English). In addition, all participants stated to have intact colour vision.

For study 1, 21 participants were recruited (twelve female, nine male; mean age and standard deviation was 26.8 ± 3.8 years, ranging from 20 to 34 years). For study 2, we recruited a fresh sample of 20 participants (ten female, ten male participants; mean and SD age 25.1 ± 5.4 years, ranging from 18 to 35 years). All participants gave written informed consent.

### Procedure

In a quiet room with dimmed illumination, participants were seated in front of a black screen (LCD monitor, 60 Hz) with a distance of approximately 70 cm. Responses were measured with temporally precise response pads (LumiTouch, Burnaby, Canada). The experimental paradigm started with two training sessions, followed by the main task. On average, one experimental session lasted between 50 and 60 minutes.

### Experimental Paradigm

#### Task

The experimental paradigm is a speeded hierarchical decision-making task with two stimulus dimensions and four response options – and will be called in the following the Speeded Inference Game – Evaluation Edition (SIG-E^2^). The game was scripted in PsychoPy (Peirce, 2007, 2008) and the open-source code can be found on GitHub (https://github.com/EvaNiessen/Speeded-Inference-Game). The goal for participants was to identify on each trial the correct target out of a set of four targets (ball, bird, ice cream, chair) based on specific rules (see below). The four targets were mapped to four spatially congruent response buttons, which should be pressed with the left and right middle and index fingers (Figure 1A). Thus, the most left target was assigned to the left middle finger, while the most right target was assigned to the right middle finger. The four targets were defined by a colour-category match that remained constant throughout the experiment. A possible target constellation is shown in Figure 1 (blue ball, green ice cream, red chair, and yellow bird).

**Figure 1.**
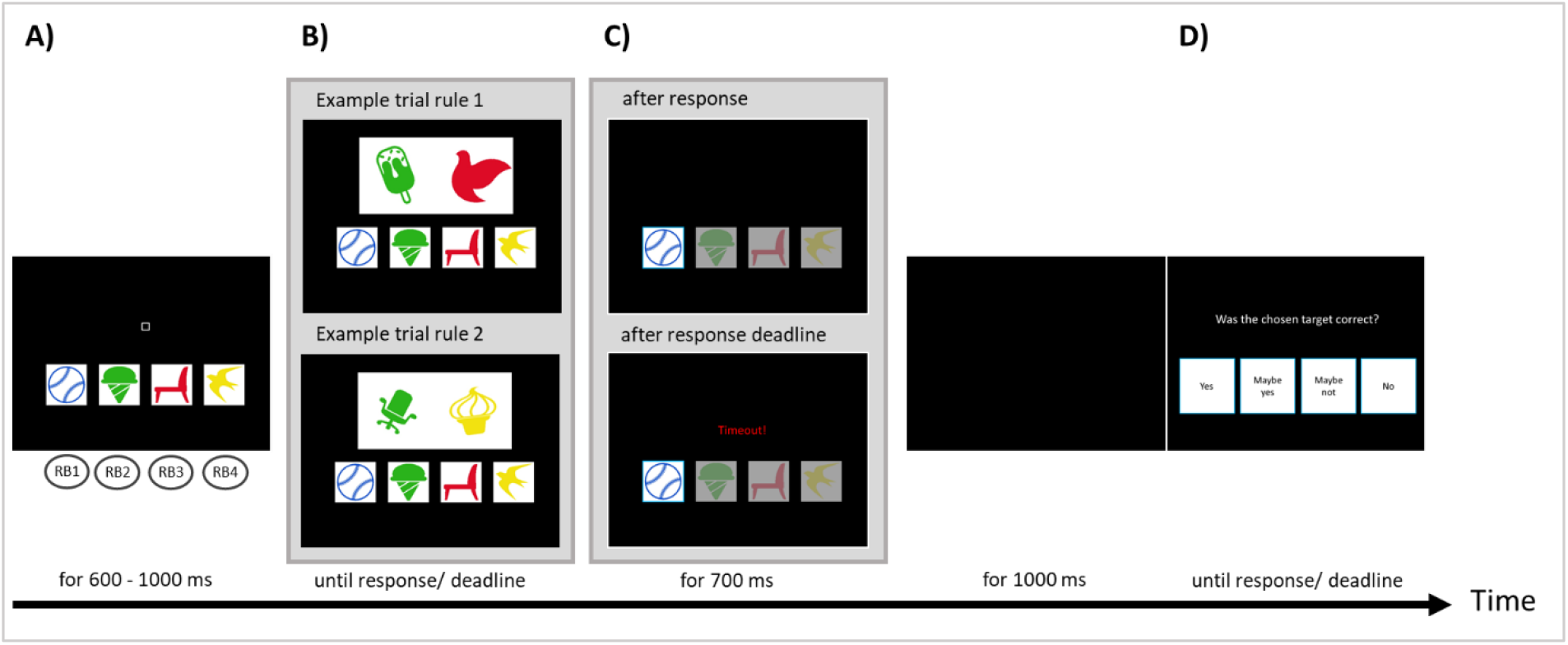
Display during the task and example trials. Only one of the displays included in B and C (inside grey background) are shown in each trial, but otherwise, the course was identical for all trials. A) Four targets are shown constantly throughout the experiment (here: a blue ball, a green ice cream, a red chair, and a yellow bird). Targets should be chosen by pressing the corresponding response button (RB; left middle finger for the blue ball, left index finger for green ice cream, etc.). B) On each trial, a stimulus appears above the targets representing two mixtures of the target dimensions. Depending on the stimuli, either rule 1 or rule 2 had to be applied. Top: Example of a trial applying rule 1, where the correct target would be the green ice cream because green ice cream is part of the stimulus. Bottom: Example of a trial applying rule 2, where the correct target would be the blue ball. One or more features of the other targets are present in the stimuli, but neither the colour blue nor a ball are present in the two stimulus items. C) Top: After a response is given by selecting one of the targets, this target is highlighted. Bottom: When no response is given until the response deadline, the word ‘Timeout’ replaces the stimulus and the software chooses a random target. The highlighting of the chosen target is identical to the top scenario and is followed by a black screen. E) An evaluation screen appears asking participants to rate the previous decision on a four-point scale using the same four RB as in the main task.

#### Rule Types^3^

A stimulus appeared above the targets on each trial and showed two items that represent a mixture of the target features (see Figure 1B, examples for the two rules). Participants were instructed to apply two rules in a hierarchical order to identify the correct target based on a given stimulus. The first rule implies that if one of the two items represents the same category *AND* the same colour of one of the targets, this target should be chosen (see Figure 1B top, example rule 1: green ice cream). If the first rule is not applicable, then the participants should apply the second rule. In that case, participants should search for the target whose category and colour are both *NOT* represented in the two items of the stimuli, i.e., the target that differed in both colour and category when compared to the two stimulus items (see Figure 1B bottom, example rule 2: blue ball). In each trial, participants had to identify the appropriate rule.

#### Performance evaluation

After a target was chosen, a 1000 ms black screen was presented; then, the participants were asked to evaluate the correctness of their decision. A rating screen appeared, asking “Was the chosen target correct?”, and participants could answer by using the same four response fingers. They could choose to answer “yes” (left middle finger) or “no” (right middle finger), and in addition, we allowed the expression of uncertainty in their evaluation and hence provided also the potential answers “maybe yes” (left index finger) or “maybe not” (right index finger, see Figure 1D).

#### Timeout

To provoke the participants’ continuous intrinsic motivation to perform well on the task, we used computer game-like characteristics. One build-in feature leading to the impression that participants were playing a game was a sudden abortion of a running trial. In these cases, before the participant responded, the stimulus was replaced by the word ‘TIMEOUT’ (written in red ink; see Figure 1C bottom). Participants were instructed to respond as fast as possible, and whenever timeouts occurred, this was a signal for too slow responses. In the case of timeouts, one of the targets was randomly chosen by the software. This randomly chosen target could be correct or incorrect, and in the next step should be evaluated by participants in an identical way as the self-chosen targets. Unbeknownst to the participants, the temporal threshold for timeouts was constantly adjusted based on the past accuracy of the participants. We implemented an adaptive algorithm to keep the level of difficulty comparable between all participants and avoid ceiling or bottom performances.

#### Adaptive algorithm

The threshold for timeouts started with 2000 ms. The threshold was adjusted after 16 trials (thus twice per block). When the average accuracy of the last 16 trials was below 60%, the threshold was increased by 10%. When the average accuracy of the last 16 trials was higher than 90%, the threshold was reduced by 10%. When accuracy lay between 60 and 90 %, the timeout threshold remained unchanged.

#### Scoring System

We used a scoring system that provided participants with an indication of their performance during the breaks between the blocks. However, we wanted to avoid trial-based feedback as behavioural adaptation should rely on internal error processing, not external feedback. Participants could gain 10 points for each correct response on the main task and further 3 points for correct evaluations. Thereby, a good performance on the main task was most important, but the evaluations also required the participants’ attention. The points for the correct evaluations were given independent of the certainty of the evaluation (e.g., the answers ‘yes’ and ‘maybe yes’ as well as ‘no’ and ‘maybe no’ were treated identically). Importantly, participants were instructed to also rate the automatically chosen target in timeout trials (at which participants could earn 3 points for that trial through correctly evaluating the randomly chosen target), which allowed us to assess the participants’ awareness for these automate decisions. During breaks, participants were presented with the maximum score that could have been reached up to that moment and the score participants had actually reached. In addition, the participants were informed about a guaranteed reward for the best participant of the current sample. Therefore, the final score of the best participant was shown in all breaks to increase the motivation of the participants to beat this high-score. The intention behind this was to prevent fatigue and boredom at the end of the experimental session and obtain optimal performance throughout the entire session.

#### Stimulus material

The two presented stimuli were drawn of a set of seven different exemplars of each target category. Thus, the participants had to respond to the concept, e.g., “ball”, and not to the specific target exemplar. This increased the task difficulty and intensified the game character of the task. The four target exemplars were constant for all participants. They were deliberately chosen because they were most representative of that category. The 28 exemplars used for the stimuli were shown to the participants before the experiment to inform the participants about the stimulus diversity (see Supplementary Material 1). The objects in the stimuli were variable in size (reduction up to 60 % possible compared to the picture’ original size) and orientation (tilted for ± 40 degrees).

#### Trial course

Figure 1 illustrates the sequence and time course of a trial. Each trial started with the presentation of the four targets and a small square representing a fixation point in the centre above the targets. After a random interval of 600-1000 ms, the stimuli appeared above the targets. The stimuli disappeared when a response was given, and for 700 ms the chosen target was highlighted (Figure 1C top) by a turquoise frame surrounding the chosen target, whereas the transparency of the three not-chosen targets was increased. This layout was identical for self-chosen targets and the automatically chosen targets after the timeout (which were additionally marked by the word ‘TIMEOUT’; Figure 1C bottom). A black screen appeared for another 1000 ms before the rating screen to evaluate the response was presented. After participants had chosen an evaluation response or no response was given after 4000 ms, the inter-trial interval (see above 600-1000 ms) started, including the presentation of the targets and the fixation square.

#### Controlling procedures

We balanced the target colour-category matching across participants, precluding an unwanted colour-based bias (e.g., the red colour could increase alertness); thus, each colour could be assigned to a different finger. The categories were always allocated to the same spatial positions.

Further, the correct target-response assignment frequency was equally distributed within each block; thus, within one block (32 trials), each of the four targets was the correct response eight times. To perfectly balance the timeout displays was not possible as the number of timeouts varied between participants. We defined a set of eight targets (each target twice), and in case of a timeout, the displayed target was drawn randomly (without replacement) from this set – thus controlling for a similar number of chosen targets independent of the correctness of that target in the given trial.

#### Practice blocks

As the target-finger mappings had to be practised to avoid errors due to difficulties in mapping targets and response fingers, we included two training sessions, which could be repeated until participants felt comfortable and the experimenter was satisfied with the performance. More specifically, participants were only allowed to continue when they showed automation in using the four response buttons (i.e., learned the target-response mapping) and when they were not frequently interrupted by timeouts (i.e., they responded as quickly as desired). The first training session was mainly intended to introduce participants to the layout of the task and familiarise them with the stimuli and targets. There was no speed limit, and trial-wise feedback should help become acquainted with the task and the two rules. The second training session resembled the main task (i.e. no feedback was given, and the timeout was used); only the threshold of the timeout was not adjusted.

#### Specifics of Study 1

Study 1 consisted of 14 experimental blocks with 32 trials (i.e., 448 trials in total). Between each block, there was a break with a self-chosen duration (minimum break was 3 seconds). Both rules had to be applied similarly often within each block, hence 50% of the time.

#### Specifics of Study 2

To investigate the impact of the frequency of the rules on error detection performance, we systematically varied the frequency of the rules. Fifteen experimental blocks with 32 trials were presented (i.e., 480 trials in total). In the first five blocks, both rules were again equally distributed (50% each rule). For the following five blocks, 75% of the trials demanded applying rule 2 and 25% of the trials the application of rule 1. Finally, this distribution was reversed for the final five blocks (75% rule 1, 25% rule 1). A constant order for every participant was used as piloting work revealed lower error rates when most trials demanded rule 1. Hence, in these blocks, the temporal threshold for the timeout was enormously shortened. With such a heavy time pressure in the (more challenging) manipulation condition, in which most trials demanded rule 2, it would have been challenging to respond before the temporal threshold was reached (i.e., the timeout occurred).

### Data and statistical analysis

#### Study 1

The response rates are first described on a descriptive level. To investigate the changes of the different response rates across the task, repeated-measures analysis of variance (RM ANOVA) were employed with the within-subject factor block (block 1 - 14) and as dependent variable either correct rate, error rate, timeout rate and error detection rate. The correct rate, error rate and timeout rate correspond to the proportion amongst all trials responded to either correctly, incorrectly or where a timeout had occurred. Detected errors were defined as those errors which were rated as being incorrect [i.e., rating question ‘Was the chosen target correct?’ answered with ‘No’ and ‘Maybe not’; (number of detected errors/ all errors)]. Undetected errors, consequently, were those errors rated as being correct (i.e., rating question ‘Was the chosen target correct?’ answered with ‘Yes’ and ‘Maybe yes’). For the influence of rule type, we conducted paired-sample t-tests for the error rate and error detection rates comparing rule 1 versus rule 2 trials [i.e., number of errors / 224 (224 is the sum of trials applying one rule type)]. Then, to assess a potential differential performance improvement due to the two rules, we conducted two 2 × 14 RM ANOVAs with the factors rule and block for the dependent variables error rate and error detection rate.

Response times (RT) were examined with a paired-sample t-test comparing RT of errors and correct responses. Changes throughout the experiment were tested with a 2 × 14 RM ANOVA with the factors accuracy and block. Finally, RT differences for rule 1 and rule 2 trials were tested with a 2 × 2 RM ANOVA with the factors rule and accuracy.

Behavioural adaptations were assessed regarding post-error slowing (PES) and post-error increase in accuracy (PIA). Paired-sample t-tests were conducted to compare PES with post-correct slowing (PCS). PES and PCS were calculated as the difference in RT of trial n+1 and trial n-1, where trial n represents the error and the correct response, respectively (Dutilh et al., 2012). Due to this approach, only trials were included if no timeout occurred before or after the error (otherwise, there would be no RT on that trial). In addition, we investigated the influence of error detection on PES by splitting errors into detected and undetected errors and compared the corresponding PES using a paired-sample t-test. The same analysis was conducted for PIA compared to the post-correct increase in accuracy (PCA).

#### Study 2

As a first step, we compared the response rates of the second study with response rates from the first study to test whether the pattern of response rates was replicable. For that, we averaged the correct rate, error rate, timeout rate, or error detection rate for the first five blocks of each experiment. In these blocks, the rule frequency was 50:50 in both studies and thus the comparison was based on the same number of trials collected under similar circumstances (e.g., time on task, habituation, fatigue). The four response rates were statistically compared with two-sample independent t-tests.

The next step was to examine whether manipulating the rule frequency led to changes in performance compared to the 50:50 rule distribution. Therefore, we used the first five blocks of the second study as a baseline condition (containing the original 50 % rule frequency) and contrasted the response rates of the baseline to the two manipulation conditions. To evaluate the influence of rule frequency, the response rates were averaged for the three parts [i.e., block 1-5 (= baseline, part 1), block 6-10 (part 2) and block 11-15 (part 3)]. Paired-sample t-tests were carried out between the baseline and part 2 and the baseline and part 3 for either correct rate, error rate, timeout rate or detection rate.

## Results and Discussion Study 1

Unless stated otherwise, all results are expressed as the mean ± standard deviation (SD). If applicable, statistical tests were Bonferroni-corrected, and the Greenhouse-Geisser correction was used when sphericity was violated. Missing values were replaced by the group average.

The mean task duration was 37.4 ± 2.8 min and varied individually because the trial durations varied with the performance (e.g. timeouts, breaks).

### Response rates

#### Descriptive statistics

As shown in Table 1, the mean error rate was 18.3 % (which in absolute terms means on average 82 error trials per person), and few trials were disrupted by a timeout (8.7 %). In 73 % of all trials, the responses were correct. In only 4 % of all ratings one of the *maybe* options were used. Therefore, for the definition of detected and undetected errors, we collapsed the ‘Yes’ and ‘Maybe yes’ ratings to represent undetected errors and the ‘Maybe not’ and ‘No’ ratings for detected errors (Stahl et al., 2020). As a result, the mean error detection rate was 69.1 %. Most automatically chosen targets after timeouts were incorrect (i.e., 73.7 ± 8.0 %). Considering the ratings of timeouts, 65.7 ± 9.8 % had been correctly evaluated (i.e., correct responses were rated as being correct and errors were rated as being incorrect) and 34.2 ± 9.8 % of the evaluations were incorrect (i.e., correct responses were rated as being incorrect and incorrect responses as being correct). Finally, most timeouts occurred on trials where rule 2 was applied (85.4 ± 6.2 %).

**Table 1.** Descriptives of task performance for studies 1 and 2.

|  | CORRECT (IN %) | ERROR (IN %) | TIMEOUT (IN %) | DETECTION (IN %) |
| --- | --- | --- | --- | --- |
| **OVERALL EXP. 1** | 73.0 ± 3.9<br>[71.2 - 74.8] | 18.3 ± 5.6<br>[15.8 - 20.9] | 8.7 ± 4.0<br>[6.9 - 10.5] | 69.1 ± 15.2<br>[62.1 - 76.0] |
| **EXP.1 PART 1**<br>**(50% RULE 1)** | 67.7 ± 4.1<br>[65.7 - 69.9] | 18.7 ± 7.6<br>[16.7 - 23.2] | 13.6 ± 6.8<br>[9.6 - 14.9] | 65.5 ± 16.5<br>[57.5 - 73.4] |
| **EXP. 2 PART 1**<br>**(50% RULE 1,**<br>**BASELINE)** | 71.0 ± 7.0<br>[67.7 - 74.3] | 16.3 ± 8.1<br>[12.5 - 20.1] | 12.7 ± 7.0<br>[9.4 - 15.9] | 63.8 ± 22.2<br>[53.5 - 74.2] |
| **EXP. 2 PART 2**<br>**(25% RULE 1)** | 73.8 ± 3.9<br>[71.9 - 75.6] | 17.6 ± 6.2<br>[14.7 - 20.5] | 8.7 ± 6.2<br>[5.8 - 11.5] | 76.3 ± 19.1<br>[67.4 - 85.2] |
| **EXP. 2 PART 3**<br>**(75% RULE 1)** | 83.4 ± 3.7<br>[81.7 - 85.1] | 12.5 ± 5.7<br>[9.9 - 15.2] | 4.0 ± 4.2<br>[2.1 - 6.0] | 63.5 ± 21.8<br>[53.3 - 73.7] |

*Note.* Group mean ± SD are shown. Square brackets include the 95% confidence interval.

#### Response rate variations across blocks

First, we tested the effectiveness of the applied adaptive algorithm, which was based on the aggregated previous accuracy of the participants. As shown in Figure 2 and tested by the RM ANOVA with the factor block, we succeeded in creating a robust and stable error rate that was not changing throughout the task [Figure 2, red line; F(13,260) = 1.002, p = .426]. Intriguingly, the detection of errors was also stable across the experiment [Figure 2, orange line; F(13,260) = 1.039, p = .407]. On the other hand, it was obvious that participants’ performance got better with time, illustrated by a significant increase in correct responses [Figure 2, green line; F(13,260) = 8.906, p < .001], which could be explained by a reduction in the occurrence of timeouts throughout the course [Figure 2, black line; F(13,260) = 17.370, p < .001].

**Figure 2.**
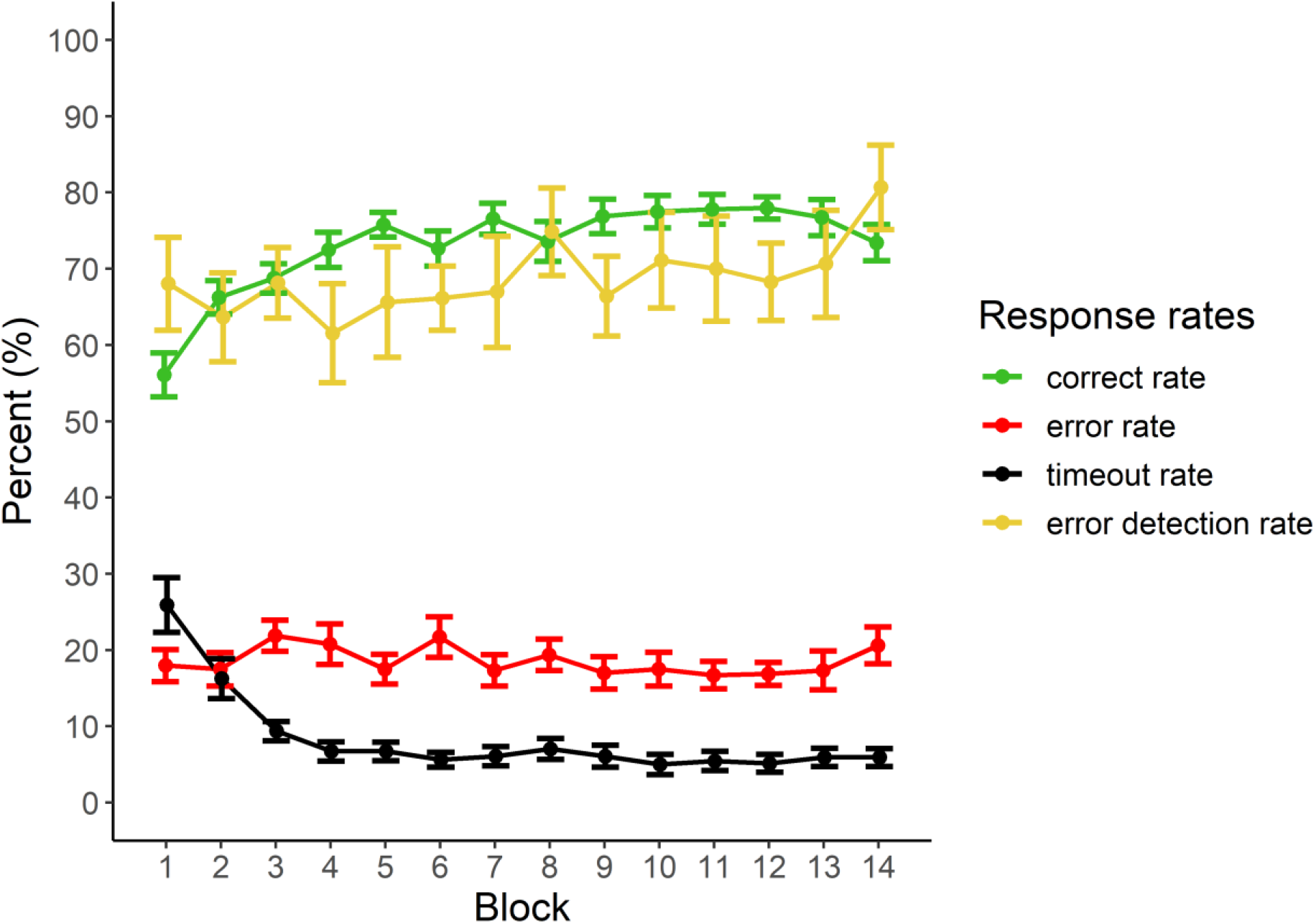
Mean ± standard error of the four response rates for each of the 14 blocks of study 1. Green line = correct rate; red line = error rate; black line = timeout rate; orange line = error detection rate.

#### Influence of rule type

We assumed that the first rule would be experienced as easier compared to the second rule. Therefore, we compared the number of errors that occurred when rule 1 should have been applied with errors occurring in trials when rule 2 had to be applied. Interestingly, the error rates did not differ significantly for the two rules [errors on rule 1 =16.3 ± 8.6 %, errors on rule 2 = 20.4 ± 9.1 %; t(20) = −1.38, p = .184]. We observed great individual difference though (see Supplementary Material 2), but these eventually balanced out on the group level. In terms of error detection, the two rules differed substantially [t(20) = −7.220, p < .001]: While most errors on rule 2 were detected (87.1 ± 16.8 %), the rating was at chance level for errors on rule 1 (48.7 ± 16.1 %). We further explored this by assessing changes in error rate and error detection rate for the two rules separately across the course of the experiment. For errors in general, the 2 × 14 RM ANOVA with the factors rule and block did not show a differential development of the two rules [non-significant interaction: F(13,260) = 1.070, p = .385]. Regarding error detection, the RM ANOVA revealed only a significant main effect of rule [due to higher error detection rates for errors on rule 2 than rule 1; F(1,20) = 61.627, p < .001)]. There was no significant main effect of block [F(13,260) = 1.561, p = .097], indicating no general increase in error detection. Finally, there was also no significant interaction [F(13,260) = 1.454, p = .135], meaning that even though the detection of errors on the second rule was generally better, this did not result in a differential change over the task (see Figure 3).

**Figure 3.**
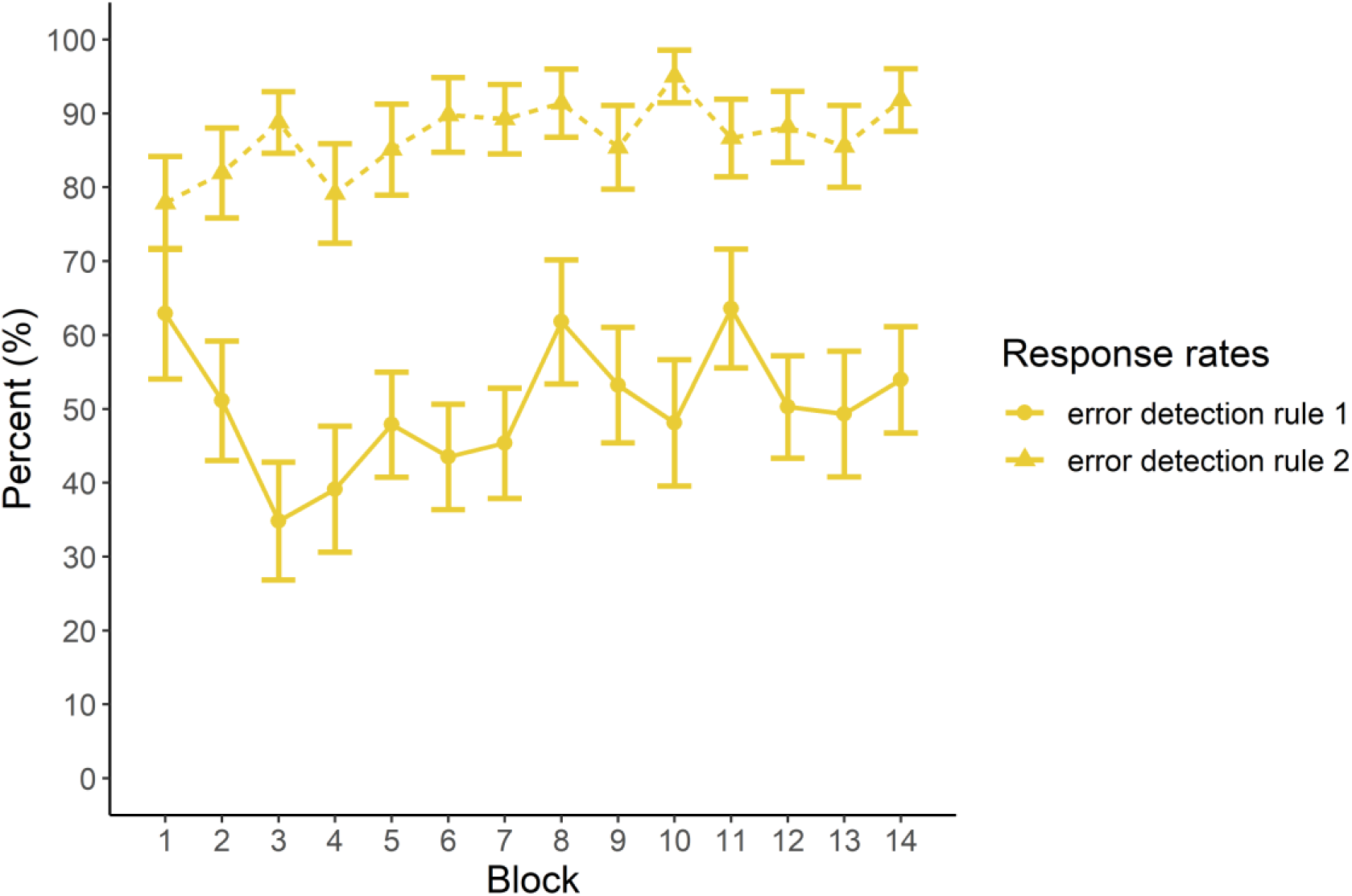
In study 1, detection rates of errors (mean ± standard error) conducted at trials demanding rule 1 (solid line) and rule 2 (dashed line) did not change across the course of the experiment. Mean detection of errors at rule 2 was better than for errors on rule 1.

### Response time

The RT of errors was significantly slower than RT of correct responses [correct = 1414.6 ± 136.1 ms; error = 1675.5 ± 164.9 ms; t(20) = −11.972, p < .001]. When inspecting the development of RTs across the experiment, the RM ANOVA revealed a main effect of accuracy [F(1,20) = 169.767, p < .001] and a significant main effect of block [F(13,260) = 10.061, p < .001]. This is in line with our interpretation above, arguing that participants became better on the task by avoiding timeouts accompanied with decreasing overall RTs. However, there was no significant interaction between block and accuracy [F(13,260) = 1.634, p = .143], indicating that the RTs became faster for both errors and correct responses.

We again were interested in the potential influence of rule type on RTs. The 2×2 RM ANOVA with the factors rule and accuracy showed a main effect of rule [F(1,20) = 237.087, p < .001] with slower RTs for rule 2 (1736.9 ± 38.3 ms) than for rule 1 (1349.2 ± 25.5 ms). As already seen, correct responses (1405.2 ± 28.3 ms) had faster RT compared to errors (1681.0 ± 33.4 ms) [F(1,20) = 321.543, p < .001]. However, the interaction between accuracy and rule was also significant [F(1,20) = 453.147, p < .001]. Post-hoc tests revealed that RTs were fastest for correct responses on rule 1 (1056.7 ± 25.4 ms), followed by errors on rule 1 (1641.8 ± 32.0 ms) and errors and correct responses on rule 2 (1753.7 ± 39.3 ms; 1720.2 ± 38.3 ms, respectively; all p < .01). This indicates that the main effect of rule type on RT was mainly driven by the difference in RT for correct and incorrect responses in trials, in which rule 1 should have been applied.

### Behavioural adjustments

A paired-sample t-test showed that PES was significantly larger (50.0 ± 78.2 ms) compared to post-correct slowing (PCS; −6.5 ± 33.4 ms) [t(20) = 2.870, p < .01]. We then investigated whether PES differed between detected and undetected errors. Differentiating errors into detected and undetected errors did, however, not result in significant differences in PES [PES detected errors = 58.2 ± 111.0 ms, PES undetected errors = 14.1 ± 114.3 ms; t(20) = 1.435, p = .167].

The same procedure was applied on post-error increases in accuracy (PIA). While we did not observe a significant difference in PIA (79.0 ± 6.9 %) compared to post-correct accuracy [PCA, 80.7 ± 5.7 %; t(20) = −1.823, p = .08], PIA significantly differed between detected and undetected errors [t(20) = 2.527, p < .02]: PIA was higher when the error was detected (80.4 ± 7.5 %) than when the error was undetected (75.6 ± 7.8 %). Thus, in terms of PIA, the awareness of an error led to greater behavioural adjustments.

### Discussion Study 1

First and foremost, we succeeded in creating an experimental task that produces a large number of undetected errors in an efficient way (i.e., in a short amount of time). The error rate was stable across the experiment, which was expected given that the adaptive algorithm was adjusted based on the past error rate. Participants were generally able to improve their performance, but this improvement was grounded in RT decreases and avoidance of timeouts – and thus not at the expense of a reduced error rate towards the end of a task. This suggests that using the past error rate for adjusting the adaptive algorithm was a successful approach. The finding of a likewise stable error detection rate was surprising. This feature of the current task, however, could be promising, because an equal distribution of detected and undetected errors throughout the experiment might reduce potential systematic biases (e.g., habituation effects at the beginning of the task or fatigue effects at the end of the task).

Further, we implemented two hierarchical rules, which both had to be applied in 50 % of the trials. This reduced surprise effects through infrequent events that are used in many standard tasks such as Go/Nogo or Flanker tasks. Despite presupposed differences in difficulty for the two rules, errors occurred similarly often. Nevertheless, the errors due to rule 1 and 2 seemed to be fundamentally different in terms of RT pattern and error awareness. While most errors due to rule 2 were detected, errors due to rule 1 were only detected in approximately 50 %, i.e., at chance level. The reasons underlying the increased number of undetected errors due to rule 1 will be further explored in study 2.

Finally, it was interesting to observe that the behavioural adjustments based on RT and accuracy did not reflect similar processes (Endrass et al., 2012): While PES was generally related to the accuracy, adjustments in PIA were more substantial when errors were detected (note that there was a trend towards significance also for the difference between PIA and PCA). While those results contradict some previous findings (Cohen et al., 2009; Kirschner et al., 2020; Klein et al., 2007), they are well in line with the adaptive orienting theory by Wessel (Wessel, 2018). According to this theory, errors first trigger a cascade of automatic processes related to the unexpected event (e.g. inhibition of ongoing cognitive processes or attentional orienting to identify the source of the error), followed by controlled processes (e.g., task-set reconfigurations) aiming to prevent errors in the future – or put differently, aiming to increase post-error accuracy. When interpreting the current findings in line with the theory, it is possible that the appearance of the rating screen in our study systematically distorted ongoing automatic processes. Consequently, larger PES was observed as a sign of a re-orienting response to the task (Notebaert et al., 2009). However, as the controlled processes probably had not been completed, an increase in cognitive control as a countermeasure to the errors’ source could not be implemented, and, therefore, we did not observe a PIA. On the other hand, error awareness, seemed to speed up the automatic and controlled processes that succeeded the incorrect response so that after detected errors, the time between the response and the rating screen was sufficiently long for the ongoing processes to be completed. Thus, the appearance of the rating screen did not disrupt or delay these processes, so that a PES was not triggered, but the adaptive measures were implemented, increasing post-error accuracy. Thus, in light of currently ongoing discussions about adaptive or maladaptive behavioural adjustments after errors, our results align with the view that PES and PIA are two distinct, but not independent measures representing maladaptive and adaptive processes, respectively. These results again highlight the critical role of error awareness, implying that the beneficial influence of error awareness should not be underestimated for processes concerning cognitive control (Endrass et al., 2012; Wessel, 2018). The here introduced SIG is a helpful tool to further investigate such processes by provoking many detected and undetected errors.

## Results and Discussion Study 2

### Behavioural results

For a replication test of the above-described findings, initial response rates of studies 1 and 2 (means of the first five blocks each) were examined to evaluate whether the second study replicated the behavioural patterns of study 1. Then, mean response rates of the baseline (i.e., mean of the first five blocks of study 2 = part 1) were compared to the mean response rates of parts 2 and 3. For brevity, we only report results of response rates here, but please see Supplementary Material 3 for results on RTs and behavioural adjustments. Please note that we refer to Table 1 for all descriptive values corresponding to the below analyses.

#### Replication

Overall, the behaviour pattern obtained in study 2 was very similar to the results obtained in study 1 and no significant group differences could be found for the four response types [correct rate p > .071; error rate p > .345; timeout rate p > .655; error detection rate p > .792]. Thus, the performance and error detection at the beginning of the two studies were similar, replicating the results of the first study.

#### Part 2 – Rule 1 applied in 25 % of trials

The paired-sample t-tests contrasting part 2 with the baseline revealed no significant differences in accuracy and the error rate (p > .175 and p > 231, respectively). The rate for timeouts was significant lower in part 2 than at baseline [t(19) = 3.214, p < .005]. However, as we saw in study 1, a reduction in the timeout rate might simply be due to habituation to the task requirements. More importantly, though, and in line with our expectations, we found a significant difference in the error detection rate between part 2 and the baseline [t(19) = −3.330, p < .004]. Thus, while the accuracy and error rate were unchanged by the differences in rule frequency, decreasing the number of trials demanding rule 1 led to a specific increase in error detection (see Figure 4).

**Figure 4.**
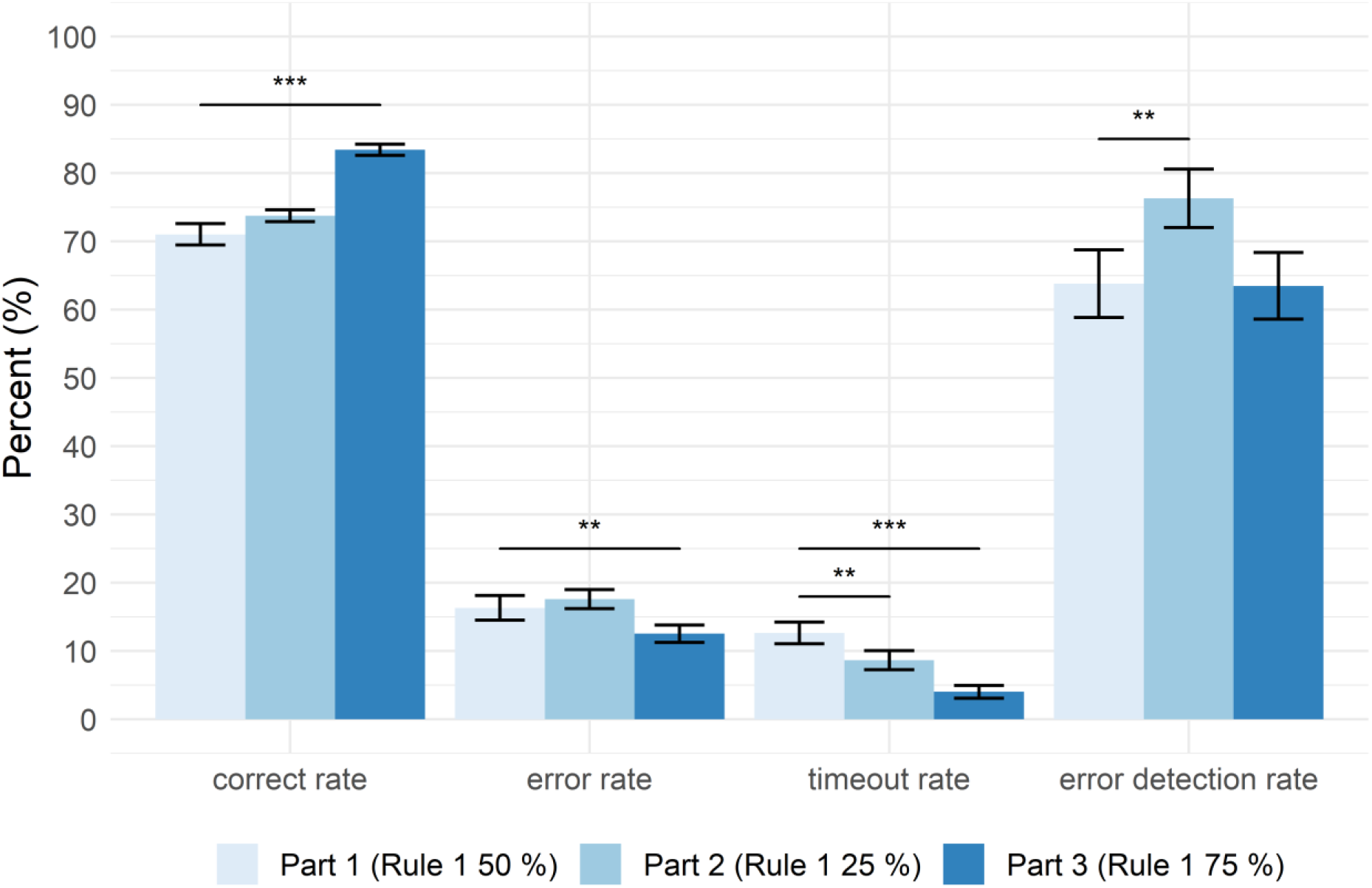
Mean ± standard error of response rates for the three parts of study 2. The correct rate, error rate, and timeout rate are represented as the percentage of all trials, whereas the error detection rate is shown as the percentage of all error trials. Asterisks indicate significant differences in comparison to part 1 (the baseline condition): * p < .05; ** p < .01; *** p < .001.

#### Part 3 – Rule 1 applied in 75 % of trials

In contrast, when adjusting rule frequency in a way that rule 1 should be applied in the majority of trials, we observed the opposite pattern of effects. Compared to baseline, the correct rate significantly increased [t(19) = −7.562, p < .001], the error rate significantly decreased [t(19) = 2.840, p < .010], and the timeout rate was also significantly lower [t(19) = 6.507, p < .001]. Crucially, the error detection rate was not affected and thus not significantly different compared to the baseline [p > .951] (see Figure 4).

### Error sources

Due to a more differentiated information stored in the updated version of the SIG used in study 2, a more detailed analysis of error sources was possible. During the practice sessions before the actual experiment, participants were trained to apply the two rules hierarchically (i.e., rule 2 can only apply if rule 1 is not applicable). However, there was no explicit cue about which rule should be used within a trial. In the second study, the distribution of errors for the two rules was again similar, replicating results from study 1 despite differences in rule frequency (note that over the whole experiment, independent of the different distributions in parts 2 and 3, both rules again occurred 50 % of trials). Averaged across all blocks of the second study, errors in study 2 occurred in 46.0 ± 13.1 % due to rule 1 and in 54.0 ± 13.1 % due to rule 2 [t(20) = −1.371, p = .186]. Importantly, we also replicated the behavioural pattern found in study 1 considering the error detection rate for both rules: While the majority of errors due to the second rule were detected (87.4 ± 16.2 %), the detection of errors due to rule 1 was at chance level (48.0 ± 21.2 %) [t(20) = −8.042, p < .001].

At a closer look at potential stimuli for rule 1, it was noticeable that there is always one target which features are not represented at all in the stimuli - which is the decision criteria for correct targets on trials demanding rule 2 (see, for example, the blue ball in Figure 1B top). This, of course, only holds if the fact that the rules are hierarchical is ignored. Thus, we wanted to assess how often participants made errors not based on a general wrong decision but based on applying the wrong rule. It turned out that 76.0 ± 10.9 % of errors at rule 1 were due to the inappropriate but eventually “correct” application of rule 2. This also explains a large number of undetected errors because of those described rule errors, only a small fraction of 36.1 ± 23.6 % was detected by participants.

### Discussion Study 2

The second study further explored the differences in error awareness for the two rules. Specifically, we wanted to test whether it is possible to manipulate the error detection rate by using different frequencies of the rule type.

When inspecting the baseline block of study 2, we found that the results of the second study convincingly replicated the behavioural pattern of study 1. The main goal of study 2 was to explore if it was possible to modulate error detection by varying the rule frequency. In study 1, we had observed that errors got less often detected in trials, where rule 1 should have been applied. Thus, we expected that by systematically increasing or decreasing the number of trials applying rule 1, we might also specifically modulate the error detection rate while keeping the error rate unchanged (because there was no difference in error rate between the two rules). Results of study 2 supported our hypotheses: We could show that error detection was significantly modulated, while at the same time, the error rate was unchanged, underlining the specificity of the manipulation. To demonstrate the effects of the manipulation, we compared behavioural changes induced by changing rule frequency in comparison to a baseline (i.e., part 1, which was identical to the design of study 1, namely a 50:50 rule distribution). The above-described successful modulations occurred in part 2, where we increased the frequency of rule 2. In contrast, increasing the frequency of rule 1 (i.e., part 3 compared to the baseline) resulted in increased correct responses and decreased errors, but did not affect the error detection rate. The SIG was initially developed to provoke many undetected errors, but as we have shown in study 2, the SIG can easily be adjusted to produce variations of specific outcome variables and for instance provoke less undetected errors (if this outcome pattern is desired). Independent of that, we observed that avoidance of timeouts seemed to reflect general learning processes throughout the task.

Finally, study 2 led to further differentiation of error types. While in study 1, it was apparent that errors due to rules 1 and 2 were fundamentally different, we further identified sources leading to (mostly undetected) errors within the category of rule 1 errors: Rule 1 errors can be differentiated based on whether rule 2 was “correctly” applied or not, i.e., whether the error was due to a misclassification about which rule should have been applied. Depending on whether or not rule 2 was applied to errors due to rule 1, the error types differed in terms of awareness and RTs (see Supplementary Material 3 for RT results). Knowing about the different error types and possibilities of provoking or countervailing them is a useful benefit of the presented task. Knowledge about the sources of errors and the easy-to-adjust open-source code of the task entails many possible applications of the game for future studies, which will be discussed in the general discussion.

## General discussion

We presented a new paradigm useful for testing error awareness. The Speeded Inference Game – Evaluation Edition (SIG-E) overcomes several weaknesses of past attempts to provoke undetected errors, such as expanding task duration or inducing perceptual uncertainty about the stimuli (cf. Kirschner et al., 2020). In addition, the paradigm has some innovative characteristics, which can easily be adjusted in future studies - as we did in study 2. One notable feature of the paradigm is the high ecological validity due to the gamification, which increases the participants’ motivation. Further, we achieved high error rates without using oddball stimuli or rare events (e.g., as in classical Go/Nogo or Flanker tasks). Only two rules needed to be memorised and autonomously applied on each trial, but when both rules occurred similarly often, errors could not be explained by surprise effects due to infrequent stimulus presentation. Another advantage, which might be especially crucial for future imaging and neurophysiological studies, is that early perceptual processes (e.g., N1 and P2 components) and decision-related processes are not overlapping in time due to the relatively long response times. A separation of perceptual and response-locked event-related potentials might hence be possible with our task. Finally, because the stimuli were shown on the screen until a response was made, we could assure that errors did not occur due to perceptual difficulties but are more likely to be related to higher-order cognitive processes (e.g., as described above for the identified rule application errors).

### Limitations

This fact, however, likewise leads to a significant limitation of the task. We created a paradigm that should be challenging and at the same time ecologically valid without relying on too many mnemonic aspects. Therefore, it was necessary to increase the complexity at several stages (e.g., four possible response buttons, implementation of category-colour matches, hierarchical rules). To counteract the complexity, we used comprehensive practise sessions before the start of the main task to ensure an internalised category-colour matching.

Another limitation was missing relevant information in the output file during study 1, which was improved in study 2 (and of course implemented in the code). Further, we had no full randomisation of the colour and category matches constituting the targets. We only randomised the colours, but kept the categories fixed on their spatial positions, as we didn’t expect those to have a significant influence on the behaviour. Still, such an influence seems possible and should be investigated in future studies.

Lastly, considering the analysis, there are two points we want to stress. First, as we saw in studies 1 and 2, errors for the two rules are different in terms of their error detection rate. Therefore, the differences between the two rules might be confounded by the underlying differences in error awareness. Second, the utility of the timeouts needs to be reassessed. While they are helpful for gamification and the implementation of the adaptive response deadline, timeout trials also introduce a new and separate response type. Participants did not respond in time, but it remains unknown at which stage in the decision-making process participants were when the timeout occurred. The subjective evaluation of timeouts is also challenging to interpret and different from evaluations after self-executed responses. Therefore, the underlying processes occurring during timeouts need further investigation.

### Future variations

As mentioned above, the paradigm was designed flexibly allowing easy modifications depending on the research question. In study 2, we already proved that changes in rule frequency could modulate error detection. If useful for the research question at hand, cueing of the current rule could be implemented to prevent too many undetected errors. Using different forms of feedback would be possible (e.g., explicit on a trial-by-trial basis or implicitly via the scoring system) if the interest of future studies lies in feedback-based learning. Further, the validity of feedback and cueing could be manipulated if interested in studying believe updating. Changing the underlying features of the timeout might pose another valuable source for interesting task variations. By increasing the volatility of the timeout, one could provoke more timeouts and hence more observed (and not committed) errors. Also, the individual difficulty level of the paradigm could be adjusted by changing the timeout threshold at the beginning of the task (which was fixed to 2000 ms in our experiments). This could potentially be individualised using RT measures obtained from the training sessions.

To conclude, we here introduced a new behavioural paradigm particularly suitable to test error awareness. It provoked many undetected errors in young adults, which so far was a challenging endeavour (Kirschner et al., 2020; Niessen et al., 2017; Rabbitt, 2002). We identified different types of errors, which could be further explored with electrophysiological measures to understand better the neural processes underlying error processing and error awareness. Considering behavioural adjustments after errors, our results support the adaptive orienting theory (Wessel, 2018) and highlight the importance of systematically gathering and integrating the subjective performance evaluation of participants – which is now possible for everyone by employing the open-source Speeded Inference Game.

## Acknowledgement

We want to thank Maren Bell for her help during the recruitment and testing of participants. A warm thank you also goes out to Dr. Alexander Geiger, who was incredibly helpful during the brainstorming processes at the beginning of this work.

## Declarations

### Ethics approval

- This study was performed in line with the principles of the Declaration of Helsinki. Approval was granted by the Ethics Committee of the Deutsche Gesellschaft für Psychologie (DGPs; reference number *EN 042018*).

### Consent to participate

- Informed consent was obtained from all individual participants included in the study.

### Consent for publication

- We did not ask participants for consent of publication, because the used and published data was fully anonymised, making a recognition of individual participants impossible.

### Availability of data and materials/ Open Practices Statement

- The datasets generated during and/or analysed during the current study are available from the corresponding author on reasonable request.

### Code availability

- The open source code for the application of the Speeded Inference Game (SIG) can be found on github (https://github.com/EvaNiessen/Speeded-Inference-Game).

## Footnotes

1 We want to clarify that when we use the term *error awareness,* we refer to the conceptual process of being conscious of the incorrect action. In contrast, when we use the term *error detection*, we refer to the operationalisation of awareness in the sense that participants volitionally reported their awareness.

2 The SIG can also be used independent of the response evaluation, representing a complex speeded decision-making task.

3 The creation of the stimuli and rules was inspired by the boardgame *Ghost Blitz* (https://de.wikipedia.org/wiki/Geistesblitz_(Spiel)).

