## Supplemental Data 1 for "Speeded Inference Game: Opening a new chapter in the assessment of error awareness"

**Supplemental Material**

1. Collection of stimuli

Used pictures in the Performance Evaluation Game, representing four different categories (balls, birds, ice creams and chairs). One picture was used as target for each category (middle picture), and seven further pictures served as stimuli (surrounding pictures). These were shown to all participants before starting the task.

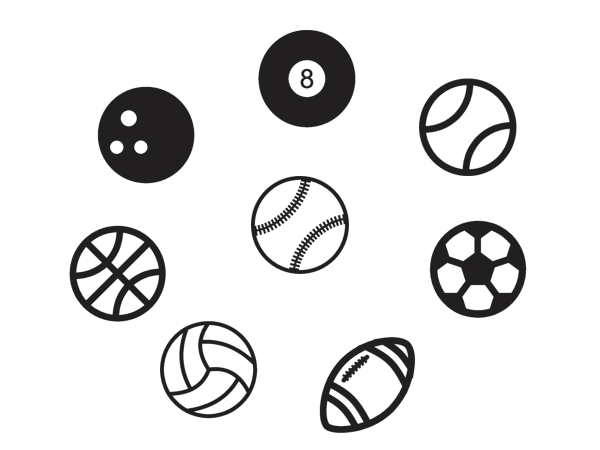

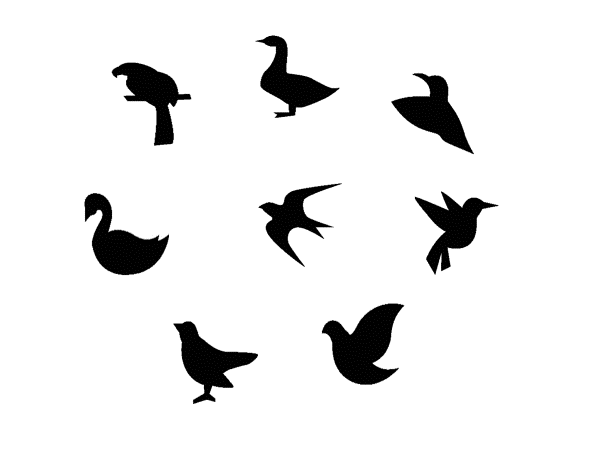

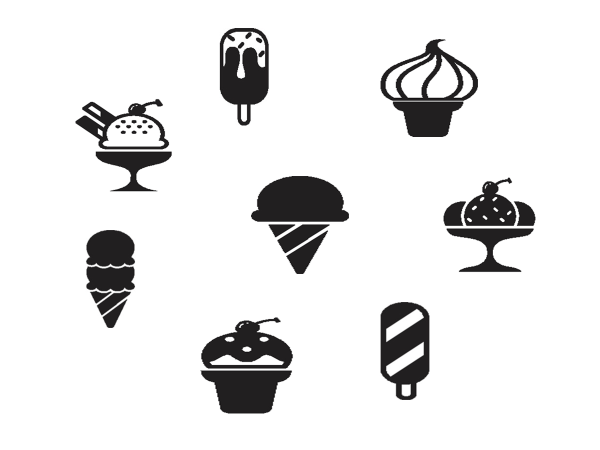

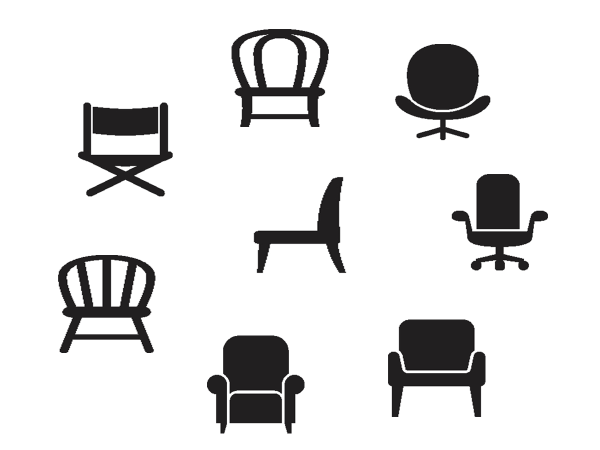

2. Individual differences of error likelihood due to the two rules

Even though on average, errors for both rules occurred similarly often (see the main manuscript), we noticed individual differences. In a post-task questionnaire, participants often reported experiencing one rule more difficult than the other. We assume that underlying strategic differences might be the reason for the individual differences (e.g., focussing on identifying a non-matching stimulus first, despite rule 1 might apply). The implementation of deliberate strategies and potential consequences on performance and error awareness should be further explored in studies employing larger samples.

The two plots below show for each participant how many percent of all errors were due to rule 1 (lighter red) and rule 2 (darker red). The dashed line at 50 % illustrates an equal distribution. Within study 1, nine participants conducted a similar number of errors due to both rules (40-60%). Nine participants experienced significant difficulties with rule 2, and hence, more than 60 % of errors where due to rule 2. Only 3 participants showed the opposite pattern and made the most errors (>60%) due to rule 1. This grouping of participants according to their error distribution looked similar for study 2 (eight participants equal errors on both rules, eight participants more errors due to rule 2, four participants more errors due to rule 1).

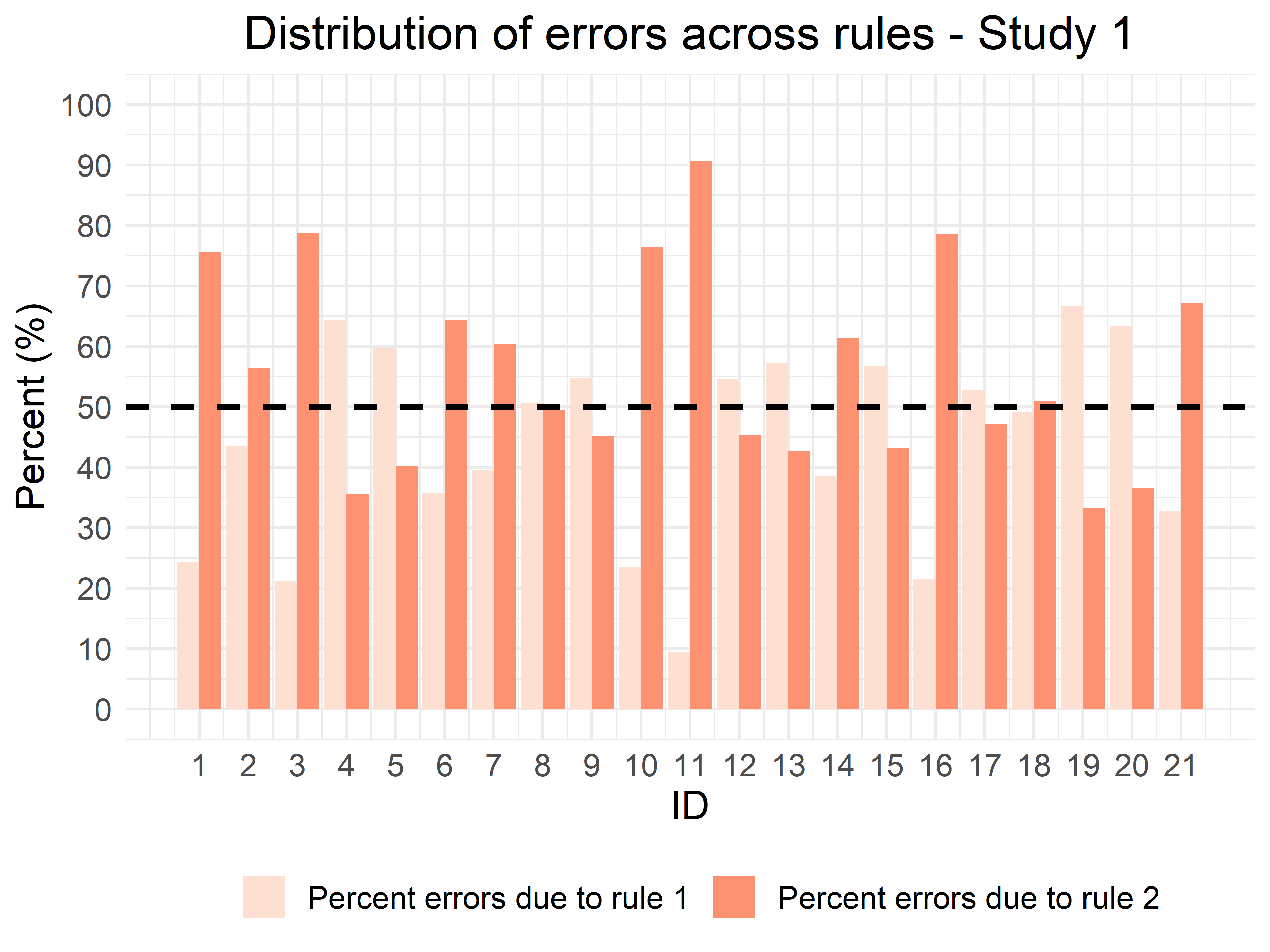

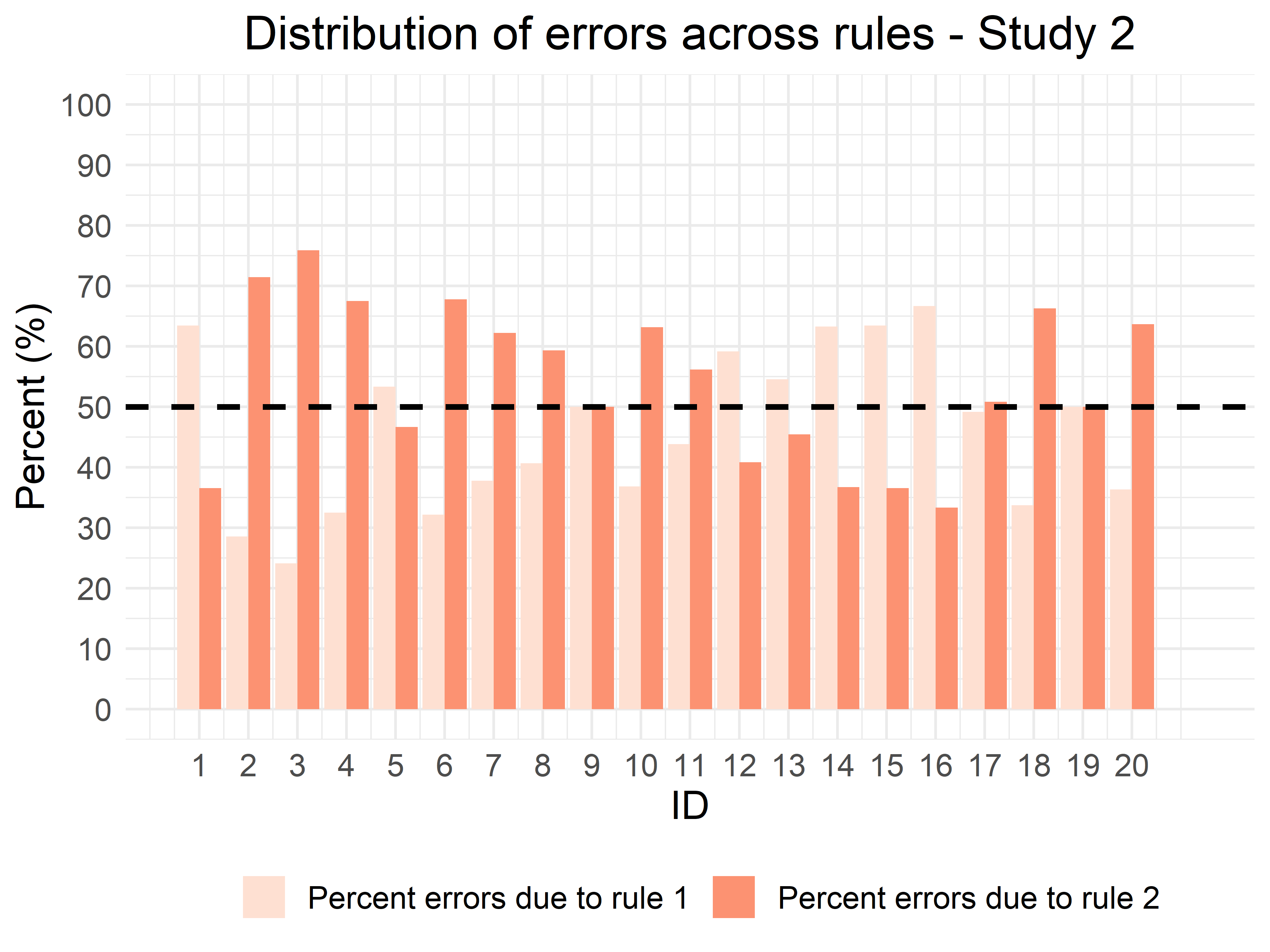

3. Additional results of study 2

*Response times*

Similar to RT results from Experiment 1, we here again saw an influence of rule type on RTs. A 2*2 RM ANOVA with the factors rule and accuracy showed a main effect of rule [F(1,19) = 295.910, p < .001] with slower RTs for rule 2 (1787.53 ± 65.29 ms) than for rule 1 (1333.81 ± 46.03 ms). In addition, correct responses (1394.49 ± 50.13 ms) had faster RT compared to errors (1726.85 ± 61.07 ms; F(1,19) = 264.473, p < .001]. However, the interaction between accuracy and rule was also significant [F(1,19) = 209.504, p < .001]. Post-hoc tests revealed that RTs were fastest for correct responses on rule 1 (998.08 ± 36.27 ms), followed by errors on rule 1 (1669.55 ± 61.20 ms) and errors and correct responses on rule 2 (1784.15 ± 64.63 ms; 1790.90 ± 67.04 ms, respectively; p > .70, all other ps < .001).

*Behavioural adjustments*

When analysing PES in a similar way as in Experiment 1 (i.e., averaged across all blocks), we did not find any differential effect as tested with one-sample t-tests. There was no significant difference between PES (24.55 ± 58.55 ms) and post-correct slowing [10.67 ± 31.21 ms; t(18) = .830, p > .418], and no significant effect of error awareness on PES [PES for detected errors = 6.47 ± 110.39 ms; PES for undetected errors = 29.00 ± 124.69 ms; t(18) = -.556, p > .585].

The same was true for behavioural adjustments in terms of accuracy. Post-error increase in accuracy (83.46 ± 8.58 %) was not significantly different from the post-correct accuracy [83.92 ± 5.76 %; t(19) = -.292, p > .773]. Again, error awareness did not influence the post-error accuracy [PIA for detected errors = 85.05 ± 9.63 %; PIA for undetected errors = 82.03 ± 9.46 %; t(19) = 1.344, p > .195].

We acknowledge that the variance in these analyses was relatively high and this could be one reason for the non-significant differences that were not expected. The manipulation of the task design possibly influenced behavioural adjustments, leading to the observed large variance across participants that putatively overshadowed any differences in PES and PIA between the three parts of study 2. Therefore, future studies with larger samples are warranted to further shed light on the role of behavioural adjustments in error awareness.

*Rule related errors – RT results*

The main text showed that errors on trials, where rule 1 should be applied, could be differentiated into two error types depending on whether or not participants applied rule 2 on these trials. Whenever participants “correctly” applied rule 2, despite the fact that the hierarchically higher rule 1 was actually to be applied, this often resulted in undetected errors. In addition, these two error types were significantly different in terms of RTs (t(19) = -2.946, p < .01): Errors were on average faster (1573.53 ± 226.67 ms) when participants did not apply rule 2 compared to when they used the incorrect rule (1705.95 ± 272.63 ms). The latter RT corresponds well to the mean RT of correct responses on trials applying rule 2 (cf. 1790.90 ± 67.04 ms taken from the second experiment).
